# Resolvin D3 controls mouse and human nociceptive functions and preclinical progression of psoriasis

**DOI:** 10.1101/715169

**Authors:** Sang Hoon Lee, Raquel Tonello, Sang-Taek Im, Hawon Jeon, Jeongsu Park, Zachary Ford, Steve Davidson, Yong Ho Kim, Chul-Kyu Park, Temugin Berta

## Abstract

Psoriasis is a chronic inflammatory skin disease caused by a complex interplay between the immune and nervous systems. Here, we found that resolvin D3 (RvD3) significantly prevented both skin inflammation and itch in a preclinical animal model of psoriasis induced by repeated applications of imiquimod. We also found that RvD3 anti-psoriatic effects resulted from the inhibition of nociceptive functions, which inhibition was replicated in human nociceptive neurons. We conclude that RvD3 and targeting nociceptive functions are promising strategies to treat psoriasis.

## Main

Psoriasis affects 2-3% of the world’s population, and is characterized by the presence of scaly skin plaques, thickened stratum corneum, infiltration and activation of inflammatory cells, arthritis and itch,.^1^ Multidirectional interactions between the nervous and immune systems have been documented in psoriasis,^2^ yet most of the emerging therapies exclusively target the latter with monoclonal antibodies against single pro-inflammatory cytokines. This approach may be limited to only a fraction of patients due to the redundancy of the immune system, be expensive over a prolonged treatment, and lead to side effects such as immunosuppression and, in some cases, even worsen the disease.^3^ Resolvins are endogenous lipid mediators generated from omega-3 polyunsaturated fatty acids with neuroimmune and pro-resolution actions^4^. Given their safety profile associated with dietary supplements, resolvins and their metabolically stable analogues may represent a novel class of therapies for the management of psoriasis. Resolvin D3 (RvD3) is a novel resolvin family member with a well-established structure and synthesis, which activates the G-protein-coupled receptor (GPCR) mouse receptor ALX/FPR2 and human receptor GPR32, and its systemic administration effectively controls leukocytes and murine arthritis.^5^ Hence, the potential therapeutic effects and mechanisms associated with administration of RvD3 in psoriasis was of great interest.

We initially investigated the potential therapeutic effects of RvD3 in a preclinical model of psoriasis consisting of repeated topical applications of imiquimod (IMQ) to murine skin (Fig 1, A), which provokes inflammatory lesions that resemble human psoriasis.^2^ IMQ application induced psoriasis-like erythema, scaliness, and lesions in mice skin (Fig 1, B), as previously described.^2^ By contrast, mice daily treated with systemic administration of RvD3 (100 ng, 1 or 2 injections daily) showed no skin lesions (Fig 1, B) and dose-dependently attenuated the development of both erythema and scaliness (Fig 1, C and D). Histologic examination of skin tissue revealed that RvD3 treatment significantly attenuated the increase in thickened stratum corneum (Fig 1, E and F), as well as the cutaneous infiltration of immune cells (Fig 1, E and G). Moreover, RvD3 also significantly decreased the transcriptional expression in skin tissues of pro-inflammatory cytokines Il17c, Il17f, and Il23a, but not Il22 (Fig 1, H), which are all associated with the activation of immune cells and pathogenesis of psoriasis.^1,2^ Taken together, these results indicate that RvD3 has potent therapeutic actions limiting psoriasiform skin inflammation, including regulating immune cells and molecules characteristic of psoriasis in mouse and human.

**Fig 1:**
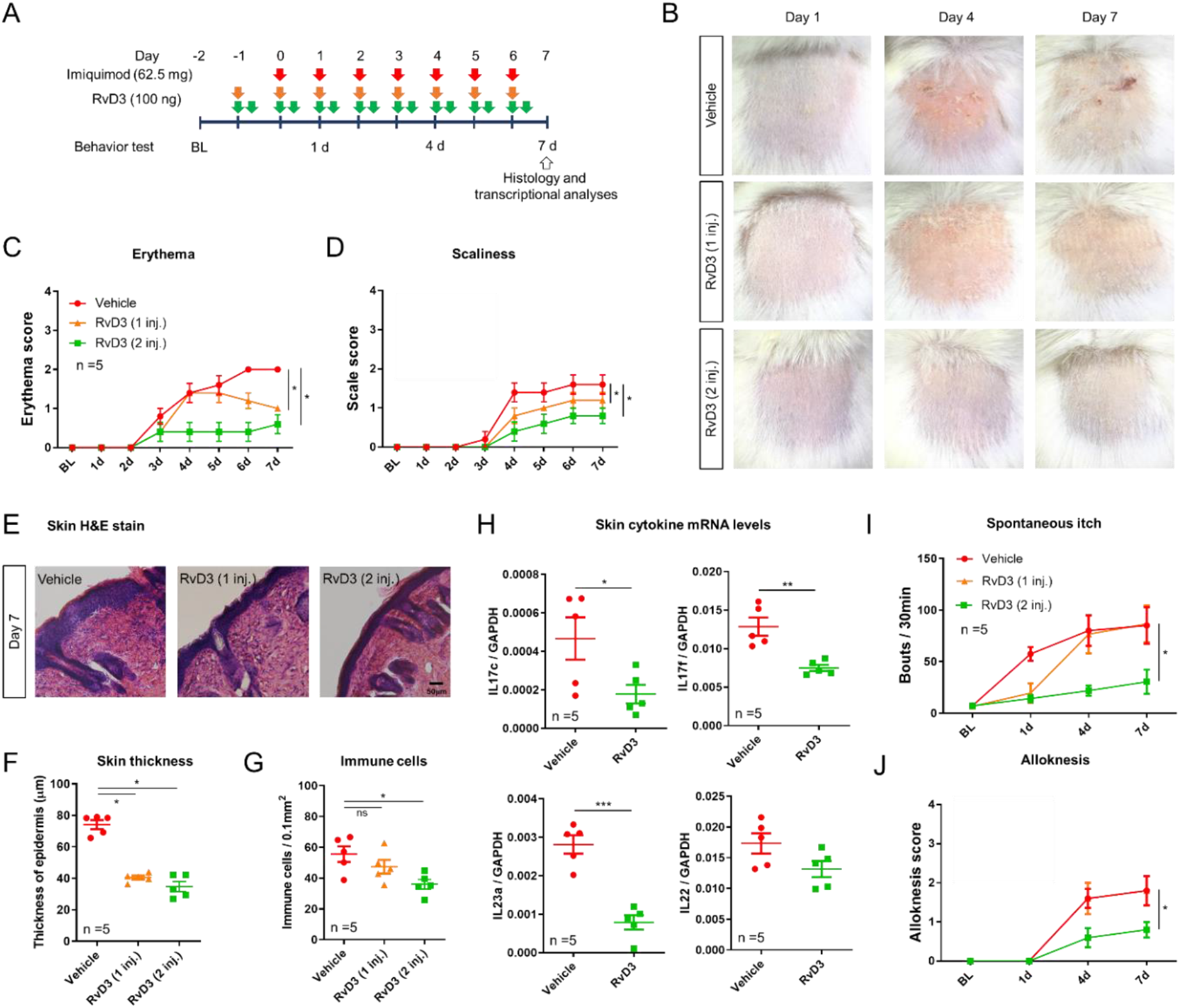
Systemic delivery of RvD3 alleviates skin inflammation and itch in a preclinical animal model of psoriasis. **A,** Timetable for drug treatment, behavioral test, and skin analysis. B, Representative images of skin lesion of psoriasis model at different time points after intraperitoneal injection of RvD3 or Vehicle. **C and D,** Repeated intraperitoneal injection of RvD3 dramatically inhibited erythema and scaliness in a dose-dependent manner. **E,** Histopathology of skin lesion from vehicle or RvD3-treated mice on day 7 of psoriasis model. Scale bar, 50μm. **F,** Dose-dependent decrease of epidermis thickness by repeated intraperitoneal injection of RvD3 on day 7 of psoriasis model. **G,** Immune cells of skin lesion were significantly reduced after daily intraperitoneal twice injections. **H,** Transcriptional expression levels of the pro-inflammatory cytokines (IL17c, IL17f, IL23a, and IL22) after repeated intraperitoneal twice injections of RvD3. Transcriptional expression levels assessed by qPCR. **I and J,** Spontaneous scratching and alloknesis of psoriasis model were significantly inhibited after daily intraperitoneal twice injections. Statistical analysis: (C, D, I, and J) Two-way ANOVA, Bonferroni post-test. (F and G) one-way ANOVA, Tukey post-tests. (H) Student’s t-test. Error bars indicate mean ± s.e.m. and ns, not significant. *p < 0.05, **p < 0.01, ***p < 0.001.

Itch is present in 60 to 90 percent of psoriasis patients,^6^ yet most preclinical studies focus on psoriasiform skin inflammation neglecting this prevalent and bothersome symptom. IMQ application induced psoriasiform spontaneous scratching and itch in response to light touch,^6^ a phenomenon known as alloknesis. RvD3 administration (2 injections daily) significantly reduced the development of counts of spontaneous scratch bouts (Fig 1, I) and alloknesis score assessed by the number of positive response to light touch stimuli (Fig 1, J). Although these experiments were designed to assess the prevention of itch in IMQ-treated mice, we also observed a reduction of spontaneous scratching in these mice immediately after the injection of RvD3. Therefore, we investigated whether a single injection of RvD3 was able to reverse established psoriatic itch 2 and 7 days after IMQ application (Fig 2, A). In agreement with a previous study,^6^ systemic administration of the histamine H1-receptor antagonist olopatadine significantly lowered spontaneous scratching 2 days, but not 7 days after IMQ application (Fig 2, B). In contrast, systemic administration of RvD3 significantly attenuated spontaneous scratching and alloknesis at both 2 and 7 days after IMQ application (Fig 2, B and see Fig S1, A), suggesting that RvD3 actions in psoriasis may act via histamine-independent mechanisms and suggests therapeutic potential for chronic itch. However, intradermal injection of RvD3 significantly lowered the number of spontaneous scratch bouts induced by cutaneous injection of histamine, but not other pruritogens such as serotonin and chloroquine (see Fig S2, A). RvD3 also did not change either motor skills assessed by rotarod, mechanical pain sensitivity assessed by graded von Frey filaments, or thermal sensitivity assessed by Hargreave’s test (see Fig S2, B, C and D).

**Fig 2:**
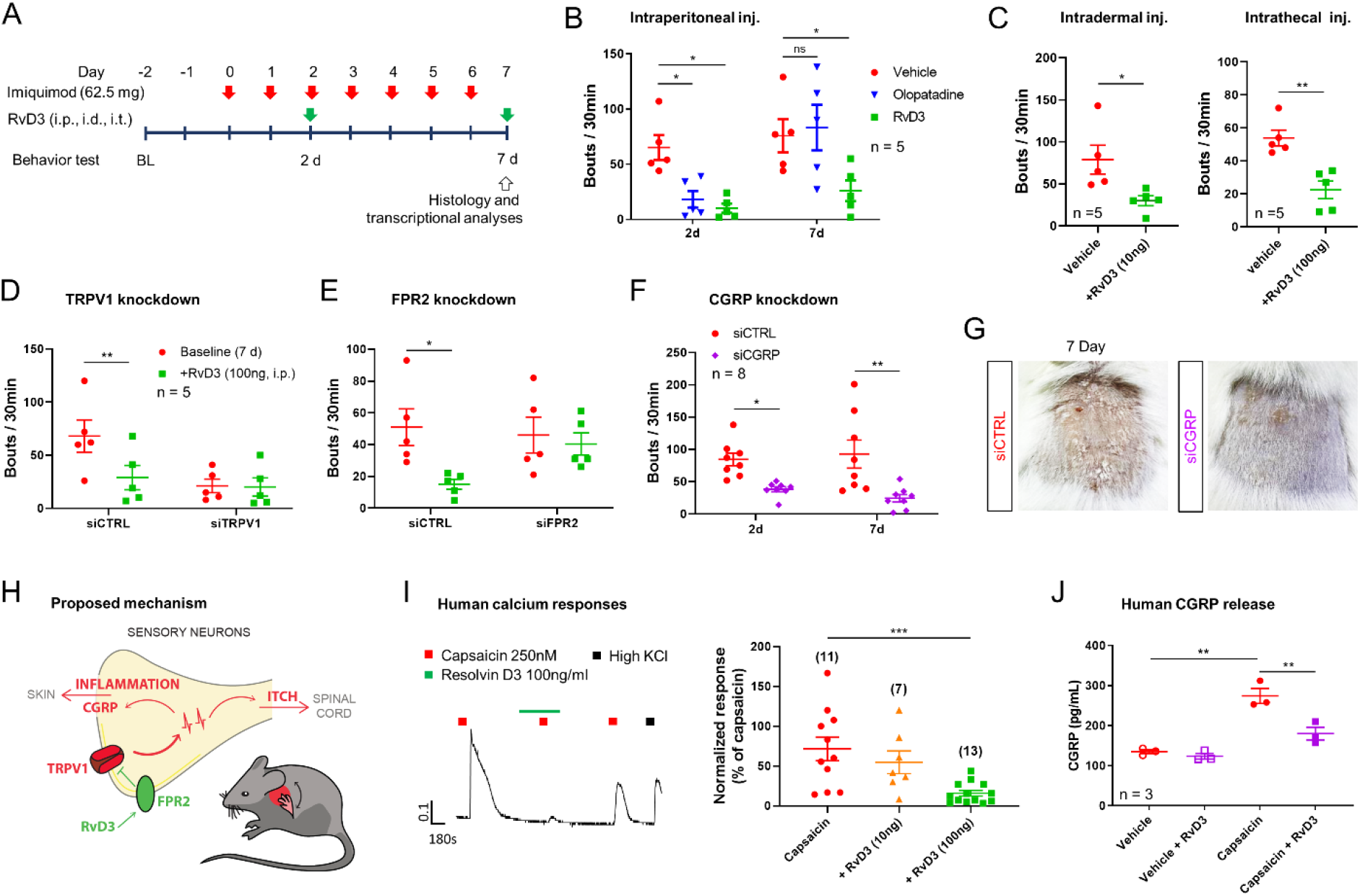
RvD3 alleviates psoriasiform skin inflammation and itch via inhibition of nociceptive functions. **A**, Timetable for drug treatment, behavioral test, and skin analysis. **B,** Spontaneous itch of psoriasis model was inhibited after Intraperitoneal single injection of RvD3. **C,** Intradermal and intrathecal single injection of RvD3 inhibited spontaneous scratching in a psoriasis model on day 7. **D and E,** Intraperitoneal single injection of RvD3 did not attenuate spontaneous scratching of TRPV1 or FPR2 siRNA-treated psoriasis model. **F,** Inhibition of Spontaneous scratching of psoriasis model after treatment with CGRP siRNA. **G,** Representative images after CGRP siRNA treatment in psoriasis model. **H,** Model of mechanism for RvD3 in sensory neurons of psoriasis model. **I,** Evoked Ca2+ responses by application of capsaicin were dose-dependently attenuated by RvD3 in human DRG neurons. **J,** Capsaicin-induced CGRP release from human DRG was inhibited by RvD3. Statistical analysis: (B, D, E, and F) Two-way ANOVA, Bonferroni post-test. (C) Student’s t-test. (I and J) one-way ANOVA, Tukey post-tests. Error bars indicate mean ± s.e.m. and ns, not significant. *p < 0.05, **p < 0.01, ***p < 0.001.

Interestingly, psoriasiform spontaneous scratching and alloknesis were decreased not only by systemic and intradermal injections but also by the intrathecal injection of RvD3 (Fig 2, C and see Fig S2, B), suggesting that RvD3 may attenuate psoriasiform itch through neuronal actions. Previous findings indicate that transient receptor potential (TRP) V1 and TRPV1–expressing nociceptive fibers control cutaneous immune responses after IMQ application.^2,7^ Pretreatment with resiniferatoxin, an ultrapotent TRPV1 agonist causing the loss of these fibers, reduced both psoriasiform itch and skin inflammation but at the same time it also abolished the anti-psoriatic effects of RvD3 (see Fig S3). Resolvins are potent inhibitors of TRPV1 and/or TRPA1^4^, and the anti-psoriatic effects of RvD3 may be mediated by its binding to the ALX/FPR2 and the inhibition of these TRP receptors in sensory neurons. Intraplantar injection of capsaicin (TRPV1 agonist) or allyl isothiocyanate (AITC, TRPA1 agonist) elicited marked nocifensive behaviors (flinching/licking) in mice. However, only capsaicin-induced nocifensive behavior was significantly reduced by RvD3 (see Fig S4, A and B). Consistently, perfusion of mouse dissociated sensory neurons with capsaicin elicited a marked TRPV1 current, and this current was dose-dependently inhibited by RvD3 (see Fig S4, B and C). Remarkably, RvD3 anti-pruritic effects were abolished in mice injected with siRNA targeting Trpv1 or Alx/Fpr2 in dorsal root ganglia (DRG) (Fig 2, D and E, and see Fig S5). To determine the involvement of GPCRs and ALX/FPR2 in DRG nociceptive neurons, we pretreated these dissociated neurons with the GPCR inhibitors pertussis toxin and GDPβS, as well as with the selective Alx/Fpr2 inhibitor PBP 10. RvD3 inhibition of capsaicin-elicited current was completely abolished by these pretreatments (see Fig S4, D, E and F). Together, these data suggest that RvD3 does not act directly on TRPV1, but its effects are mediated by the G-protein-coupled receptor ALX/FPR2 in DRG nociceptive neurons.

It is intriguing that RvD3 neuronal actions not only reduced psoriasiform itch but also skin inflammation. Given that TRPV1–expressing nociceptive fibers release numerous neuropeptides associated with itch and inflammation in the periphery^8^, we explored the effect of RvD3 on their transcriptional expression levels and found that RvD3 significantly attenuated the transcriptional expression levels of the calcitonin gene-related peptide (CGRP) in DRG of mice treated with IMQ (see Fig. E6, A). To determine whether nociceptive neurons released CGRP in skin tissue and the effects of IMQ and RvD3, we used an *ex vivo* organ culture of skin explants (see Fig S6, B). A robust increase in CGRP release was observed from skin biopsies collected 7 days after IMQ treatment, which was significantly attenuated by the co-treatment with RvD3 (see Fig S6, B). Similarly, capsaicin elicited increases in the transcription and release of CGRP in dissociated DRG neurons, which were significantly reduced by co-treatment with RvD3 (see Fig S6, C), suggesting CGRP as major mediator of psoriasiform itch and skin inflammation. Although a single systemic treatment with a CGRP antagonist did not alter spontaneous itch after IMQ (data not shown), long-lasting downregulation of CGRP expression in DRG neurons (see Fig S7, A) resulted in significant attenuation of both psoriasiform itch (Fig 2, F, and see Fig S7, B), as well as skin inflammation as indicated by the reduction of erythema, scaliness, thickened stratum corneum, and infiltration of immune cells (Fig 2, G, and see Fig S7, C-G).

Collectively these findings indicate that nociceptive neurons and their signaling have a major impact in a preclinical model of psoriasis, and reveal the molecular mechanisms by which RvD3 via FPR2, TRPV1 and CGRP attenuates both psoriasiform itch and skin inflammation (Fig 2, H). Interestingly, small-scale human clinical trials have shown the effectiveness of local neuronal treatment reducing CGRP in improving plaque psoriasis.^9^ Are our findings relevant for a clinical treatment of psoriasis? Interestingly, human DRG tissue expresses GPR32 (Fig S8) suggesting that RvD3 may have a similar role in blocking human TRPV1 receptor and CGRP release. Therefore, we tested whether RvD3 can block TRPV1 function and CGRP in human DRG neurons obtained from non-diseased donors. Calcium imaging of nociceptive human DRG neurons showed that capsaicin-induced responses were significantly reduced by RvD3 treatment (100 ng/ml, Fig 2, I). Furthermore, capsaicin-induced release of CGRP was also reduced by RvD3 treatment (Fig 2, J). In conclusion, RvD3 controls mouse and human nociceptive functions and could provide a novel therapeutic approach in psoriasis.

## Acknowledgments

The project was supported by grant from NIH (AR068012 to S.D., NS106264 and NS113243 to T.B.) and by grants from the National Research Foundation of Korea (NRF-2017M3C7A1025600 to C.-K.P.).

## Author contributions

S.H.L. conducted behavioral tests, immunohistochemistry, electrophysiology and biochemical experiments, as well as participated in experimental design and interpretation. R.T. contributed to the behavioral data and analysis. S-T.I., H.J. and J.P. carried out the electrophysiological experiments. Z.F. and S.D. performed the calcium imaging experiments. S.D., Y.H.K., C-K.P. critically assisted part of the project and revised the manuscript. S.H.L and T.B. conceived and supervised the project, as well as wrote the manuscript.

## Supplementary Materials and Methods

### Reagents

We purchased resolvin D3 (cat. no. 13834) from Cayman Chemical, 5% imiquimod cream from Perrigo (Dublin, Ireland), selective and cell permeable FPR2 antagonist (PBP 10, cat. no. 4611) from Tocris (Minneapolis, MN), resiniferatoxin (cat. no. R8756), capsaicin (cat. no. M2028), AITC (cat. no. 377430), histamine (cat. no. H7125), chloroquine (cat. no. C6628), serotonin (cat. no. H9523), and olopatadine hydrochloride (cat. no. O0391) from Sigma (St Louis, MO), Mouse Fpr2-targeting siRNA (Assay ID s66214), CGRP-targeting siRNA (Assay ID s233632), TRPV1-targeting siRNA (Assay ID s101403), and non-targeting siRNA (cat. no. 4390844) were purchased from Thermo Fisher Scientific (Waltham, MA). In vivo-jetPEI^®^ (cat. no. 201-10G) by Polyplus (New York, NY) was mixed with siRNA to increase the uptake of siRNA by DRGs.

### Animals

All protocols for animal experiments were approved by the Institutional Animal Care & Use Committee of the University of Cincinnati according to the National Institutes of Health Guide for the Care and Use of Laboratory Animals. Adult CD-1 mice (males, 8–10 weeks, Charles River Laboratories, Wilmington, MA) were used for behavioral and biochemical studies. Young CD-1 mice (6 weeks) were used for electrophysiological studies in dissociated neurons from dorsal root ganglia (DRGs). All animals were housed under a 12-h light/dark cycle with food and water available ad libitum. Sample sizes were estimated based on our previous studies for similar types of behavioral, biochemical and electrophysiological analyses.^10–12^

### Imiquimod induced psoriasis animal model

A psoriasis animal model of psoriasis was generated as previously described.^6^ Briefly, fur on the rostral back was trimmed with electric clippers and then removed with an electric shaver. The mice received a daily topical application of 62.5 mg 5% imiquimod cream on the shaved back skin (2 cm × 2 cm) for seven consecutive days. Control mice were treated similarly with a control vehicle cream (Vaseline, Trumbull, CT).

### Evaluation of skin inflammation severity

To score the severity of the nape skin, an objective scoring system was followed previous research paper based on the Psoriasis Area and Severity Index (PASI). Erythema and scaliness were scored independently on a size from 0 to 4: 0, absent; 1, mild; 2, moderate; 3, severe; 4, very severe. The level of erythema and scaliness were blindly scored by trained experimenter using a scoring table with red taints and dead skin cells.^13–15^

### Denervation

To examine the roles of TRPV1-expressing C-fibers in psoriasis, mice were injected once with capsaicin analogue resiniferatoxin (RTX, 50 μg/kg, subcutaneously) seven days prior the imiquimod treatment, as described previously.^16^ Although denervated mice exhibited insensitivity to noxious heat stimuli, overall behavior qualitatively remained unaltered in RTX-treated mice.

### Behavioral testing

Animals were habituated to the behavioral testing apparatus for 1h on each of the two days preceding data collection. All experimenters were blinded to the treatment/condition.

#### Spontaneous itch

To quantify scratching induced in absence of any experimental stimulus (spontaneous itch), mice were placed in chambers (15 × 25 × 10 cm) and video recorded for a period of 30 min; bouts of hindlimb scratching were counted offline.^10^

#### Nape stimulation assay (Alloknesis)

To quantify itch-related scratching induced by mechanical stimulation, mice were placed in chambers and a 0.16 g von Frey hair was applied to the nape for 3 s to 5 randomly selected sites of around the cream application area.^6^ The alloknesis score was the total number (0-5) of hindlimb scratching responses directed to the site of mechanical stimulation.

#### von Frey assay

To assess the sensitivity of the glabrous skin to light punctate mechanical stimulation, mice were placed in a plastic chamber on an elevated wire grid and the lateral plantar surface of the hindpaw was stimulated with calibrated von Frey monofilaments (0.02-1.4 g; Stoelting Co., Wood Dale, IL). The 50% paw withdrawal threshold for the von Frey assay was determined by the up-down method^17^.

#### Hargreaves test

To measure radiant heat pain, mice were placed in a plastic chamber and the plantar hindpaw surface was exposed to a beam of radiant heat (IITC Life Science Inc., Woodland Hills, CA). The latency to paw withdrawal was determined in three trials per hindpaw and averaged per animal, with a 10 min interval between trials.^11^ A cutoff time of 20 s was set to prevent tissue damage.

#### Motor function testing

To measure motor function, mice were trained for 5 min at a constant speed of 4 rpm on the rotarod (IITC Life Science Inc.). Mice were tested three times separated by 10 min intervals. During the tests, the speed of rotation was accelerated from 5 to 30 rpm in 3 min. The falling latency was recorded and averaged.^18^

#### Chemical nociception tests

To assess pain induced by chemical agents, a 30-gauge needle was used to inject either capsaicin (200 pmole / 20 μl) and AITC (200 nmole / 20 μl) subcutaneously into the plantar hindpaw. The time spent licking, flinching, and biting the injected hindpaw was recorded for 5 min.^19^

#### Chemical itch tests

The pruritogens chloroquine (100 μg; Sigma), histamine (100 μg; Sigma), and serotonin (4 μg; Sigma) were dissolved in 0.9% sterile saline and injected intradermally in the cheek in a volume of 20 μl. The behavior of each animal was video-recorded over the following 30 min, and the number of hindpaw scratch bouts was counted.^10,20^

### *In vivo* knock-down

Selective siRNA and non-targeting control siRNA were synthesized by Thermo Fisher Scientific. siRNA was dissolved in RNase-free water at the concentration of 1.4 μg/μl as stock solution, and mixed with In vivo-jetPEI^®^, 10 min before injection, to increase cell membrane penetration and reduce the degradation. In vivo-jetPEI^®^ was dissolved in 5% glucose, and 1 μg of siRNA was mixed with 2.62 μl of In vivo-jetPEI^®^. We intrathecally injected 10 μl of siRNA (3 μg) mixture once a day to knockdown gene expression.^11^

### Immunohistochemistry

Mice were anesthetized terminally with isoflurane and perfused through the ascending aorta with saline, followed by 4% paraformaldehyde. DRGs and spinal cords were removed and post-fixed in the same fixative overnight. DRG sections (12 μm) were cut in a cryostat and processed for immunofluorescence. Tissue sections were blocked with BlockAid™ blocking solution (cat. no. B10710, Thermo Fisher Scientific) for 30 minutes and incubated overnight at 4°C with the following primary antibodies against calcitonin gene-related peptide (CGRP; goat, 1:500, cat. no. ab36001, Abcam), TRPV1 (Guinea pig, 1:1000, cat. no. NB300-122, Novus biologicals) and 49-6-diamidino-2-phenylindole dihydrochloride (DAPI; 300 nmol/L, cat. no. D1306, Thermo Fisher Scientific). Sections were then incubated for 1 hour at room temperature with Alexa Fluor 594 (1:500, cat. no. A-11076, Thermo Fisher Scientific) and 555 (1:500, cat. no. A-21432, Thermo Fisher Scientific) conjugated secondary antibodies. Immunostained tissues were then examined under an Olympus fluorescence microscope (BX63), and images were captured with a high resolution CCD Spot camera (cat. no. DP80, Olympus) and analyzed with CellSens (Olympus). All image capture and intensity quantification were performed comparing samples from all experimental groups, prepared with the same staining solutions, then measured using identical display parameters. For immunoreactive intensity quantification, intensity was quantified by individuals who are blinded for the genotype using 3 sections per mouse and 5 mice per group.^21^

### Western blot

Protein samples were prepared in the same way as pervious study.^10^ Samples were mixed with Ripa lysis buffer (cat. no. 20-188, Milliporesigma, Burlington, MA) and kept in a cryogenic freezer and samples were homogenized using a tissue homogenizer and total protein was extracted. Concentration of protein were measured by using the Qubit protein assay kit (cat. no. Q33211, Thermo Fisher Scientific). Twenty micrograms of proteins were loaded for each lane and separated by NuPAGE™ 4-12% Bis-Tris Protein Gel (cat. No. NP0323BOX, Invitrogen). Western blotting membranes were blocked with 2% BSA for 1 hr and incubated for overnight at 4°C with FPR2 (rabbit, 1:1000, cat. no. NLS1878, Thermo Fisher Scientific), and GAPDH (mouse, 1:5000, cat. no. MA5-15738-HRP, Thermo Fisher Scientific). Samples were incubated at rabbit (1:2000, cat. no. 7074S, cell signaling) secondary antibody and developed in SuperSignal™ West Dura Extended Duration Substrate (Thermo Fisher Scientific). The membranes were scanned by iBright™ FL1000 Imaging System (Thermo Fisher Scientific) and specific bands were evaluated by apparent molecular sizes. The ratio of band intensities of protein normalized to GAPDH was calculated with Image J software.

### Whole-cell patch clamp recordings in cultured mouse DRG neurons

DRGs from all spinal levels of 6-8 week old mice were removed aseptically and incubated with collagenase (5 mg/ml, Roche) / dispase-II (1 mg/ml, Roche) at 37°C for 40 min, then digested with 2.5% trypsin (Invitrogen) for 7 min at 37°C, followed by 0.25% trypsin inhibitor (Sigma). Cells were mechanically dissociated with a flame polished Pasteur pipette in the presence of 0.05% DNAse I (Sigma). DRG cells were plated glass cover slips and then plated onto glass coverslips previously coated with a solution of 0.1 mg/ml poly-L-ornithine. DRG cells were grown in a neurobasal defined medium (with 2% B27 supplement, Invitrogen), at 37 °C, with 5% CO2. DRG neurons were grown for 18 hours before use.^22,23^ Whole-cell current-clamp recordings were performed at room temperature to measure currents with HEKA EPC10 (HEKA). The patch pipettes were pulled from borosilicate capillaries (Chase Scientific Glass Inc., Rockwood, CA, USA). When filled with the pipette solution, the resistance of the pipettes was 4 ~ 6 MΩ. The recording chamber was continuously perfused (2-3 ml/min). Series resistance was compensated for (> 80%), and leak subtraction was performed. The Pulse v8.30 software (HEKA) was used during experiments and analysis. The internal pipette solution was composed of (in mM): 140 KCl, 1 CaCl_2_, 2 MgCl_2_, 10 EGTA, 10 D-glucose and 10 HEPES adjusted to pH 7.3 with NaOH, osmolarity 295 - 300 mOsm. Extracellular solution contained (in mM): 140 NaCl, 5 KCl, 2 CaCl_2_, 1 MgCl_2_, 10 HEPES, 10 D-glucose, adjusted to pH 7.3 with NaOH, osmolarity 300-310 mOsm. Voltage-clamp experiments were performed at a holding potential of −60 mV.^10,22,23^

### Ca2^+^ imaging of cultured human DRG neurons

Human L4 and L5 DRG were recovered from consented organ donors in collaboration with LifeCenter, Cincinnati and the University of Cincinnati Medical Center and prepared for calcium imaging as described.^24^ Briefly, hDRG neurons were incubated with Fura-2-AM (3 μg/mL; ThermoFisher) for 40 min at 37°C in Neurobasal-A Media with B27, 2mM Glutamax, 5% FBS, and 1% penicillin streptomycin (Gibco). Extracellular solution contained (in mM) 130 NaCl, 5 KCl, 10 HEPES, 2 CaCl_2_, 1 MgCl_2_ and 30 D-(+)-glucose, pH 7.3. Neurons were illuminated through a 10X immersion objective on an Olympus BX51 microscope by 365 nm and 385 nm wavelength LEDs using a pE-4000 (CoolLED). Images were acquired on a Rolera Bolt CMOS camera connected to a PC running Metafluor software (Molecular Devices). Neuronal identity was confirmed at the end of each experiment with a response to 50mM KCl solution. All drugs were applied via bath perfusion at a flow rate of ~2 mL/min.

### Primary mouse DRG neurons culture for molecular biology experiments

DRGs from all spinal levels of 8 weeks old mice removed aseptically and incubated with Papain (Sigma, cat. no. P3125) at 37°C for 20 min, then incubated with Collagenase (Sigma, cat. no. C6885) for 20 min at 37°C. Cells were mechanically dissociated with a flame polished Pasteur pipette in DMEM. DRG cells were plated 8 wells glass slide and then plated onto well previously coated with Cell-Tak (Corning, cat. no. 354240) for qPCR and ELISA. DRG cells were grown in DMEM medium (with 10% FBS, 1% penicillin and streptomycin, 1 mM 5-fluoro-2-deoxyuridine, 1 mM uridine, and 25 ng/ml nerve growth factor), at 37 °C, with 5% CO2. DRG neurons were grown for 24 hours before use.

### Enzyme immunoassay (EIA)

CGRP content of culture medium was determined by EIA according to the manufacturer’s instructions (cat. no. 589001 for mouse, 589101 for human, Cayman Chemical, Ann Arbor, MI). Briefly, Medium added in duplicate to coated wells along with a CGRP standard series and incubated at 4 °C for overnight. Plates were washed thoroughly before addition of Ellman’s Regent. Ellman’s reagent was applied and absorbance (405 nm or 414 nm) read on a microplate spectrophotometer (Wallac Envision, PerkinElmer, Waltham, MA).

### CGRP release assay from skin explants

Skin punch biopsies (10 mm) were collected from the nape of psoriasis model, and rapidly transferred to 24-well plates containing 1 mL of DMEM. Explants were incubated at 38°C for 30 min. After incubation, the bath supernatant from the organ cultures was collected, and assayed to measure CGRP concentration with the CGRP EIA (cat. no. 589001, Cayman Chemical) according to manufacturer’s instructions.^25^

### Quantitative real-time RT-PCR (qPCR)

We performed qPCR in the same way as previous study.^21^ Mice were terminally anesthetized with isoflurane and cervical DRGs and skin of nape were rapidly removed treatments. Total RNA was isolated using a Drirect-zol (cat. no. R2072, Zymo research, Irvine, CA) and quantified from Nanodrop readings (Simplinano™, GE Healthcare, Chicago, IL). cDNA was generated using a High Capacity RNA-to cDNA Kit (Applied Biosystems, Carlsbad, CA). The sequences of primers are listed in Table 1. We performed gene-specific mRNA analyses using the QuantStudio 3 Real-Time PCR system (Thermo Fisher Scientific, Waltham, MA). Quantitative PCR amplification reactions contained the same amount of reverse transcription product: 5 μl of PowerUp™ SYBR^®^ Green Master Mix (Applied Biosystems, Carlsbad, CA) and 1 μl of 1.5 μM forward and reverse primers in a final volume of 10 μl. The thermal cycling conditions were: 3 min of polymerase activation at 95 °C, 45 cycles of denaturation at 95 °C for 10 s, and annealing and extension at 60 °C for 30 s, followed by a DNA melting curve for the determination of amplicon specificity. The expression level of the target mRNA was normalized to that of GAPDH mRNA and analyzed using the standard 2^-ΔΔCT^ method.

### Reverse-transcription polymerase chain reaction (RT-PCR)

cDNA was synthesized in the same way as qPCR. Samples were diluted 2:100 and used as template for PCR experiments. The following primer pairs were used: mouse Fpr2 (forward, 5’-ACTGTGAGCCTGGCTAGGAA-3’; reverse, 5’-CATCAGTTTGAGCCCAGGAT-3’), mouse Gapdh (forward, 5’-TGAAGGTCGGTGTGAACGAATT-3’; reverse, 5’-GCTTTCTCCATGGTGGTGAAGA-3’), human GPR32 (forward, 5’-TTTGCCAGTAACTGCCTCCT-3’; reverse, 5’-TGTCAGAGTTGAACGCCAAG-3’), human Gapdh (forward, 5’-ACCCAGAAGACTGTGGATGG-3’; reverse, 5’-TTCTAGACGGCAGGTCAGGT-3’).

### H&E staining

Mice were euthanized by CO2 inhalation and intracardially perfused with 20 mL of ice-cold PBS, followed by 20 mL of PBS/4% para-formaldehyde (PFA, cat. no. 19210, Electron Microscopy Science, Hatfield, PA). Psoriatic skin lesion samples were dissected, post-fixed for 12 h at 4 °C in PBS/4% paraformaldehyde solution, embedded in tissue freezing medium (Electron Microscopy Sciences), cut into 30μm-thick sections, and stained using hematoxylin and eosin (H&E) by the University of Cincinnati Histopathology Core. Stained sections were imaged by light microscopy on Olympus fluorescence microscope (BX63), and images analyzed with CellSens (Olympus). The number of immune cells and thickness of epidermis was quantified by individuals who are blinded for the genotype using 4-5 sections per mouse and 4-5 mice per group.^26^

### Statistical analysis

Values are reported as the mean ± s.e.m. For comparison between two groups, Student’s t-test was used. For single-point comparison between >2 groups, One-way ANOVA followed by a Tukey post-hoc test was used. For the time course comparison between >2 groups, 2way ANOVA for multivariate linear models was used. Significance was labeled as: ns, not significant, p ≥ 0.05; *p < 0.05; **p < 0.01; ***p < 0.001.

**Suppl. Table 1.**
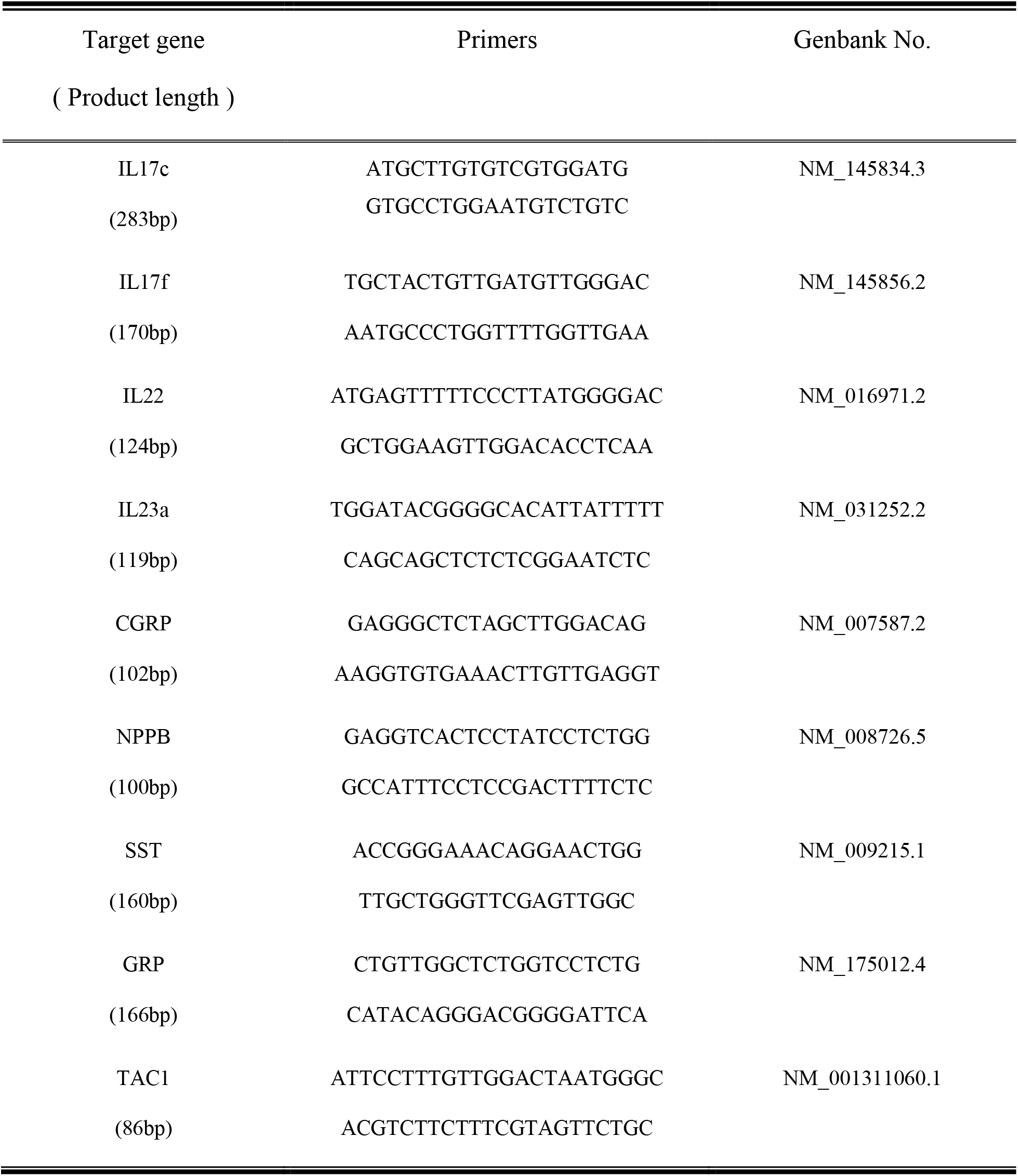
Primer sequences for qPCR

**Fig S1:**
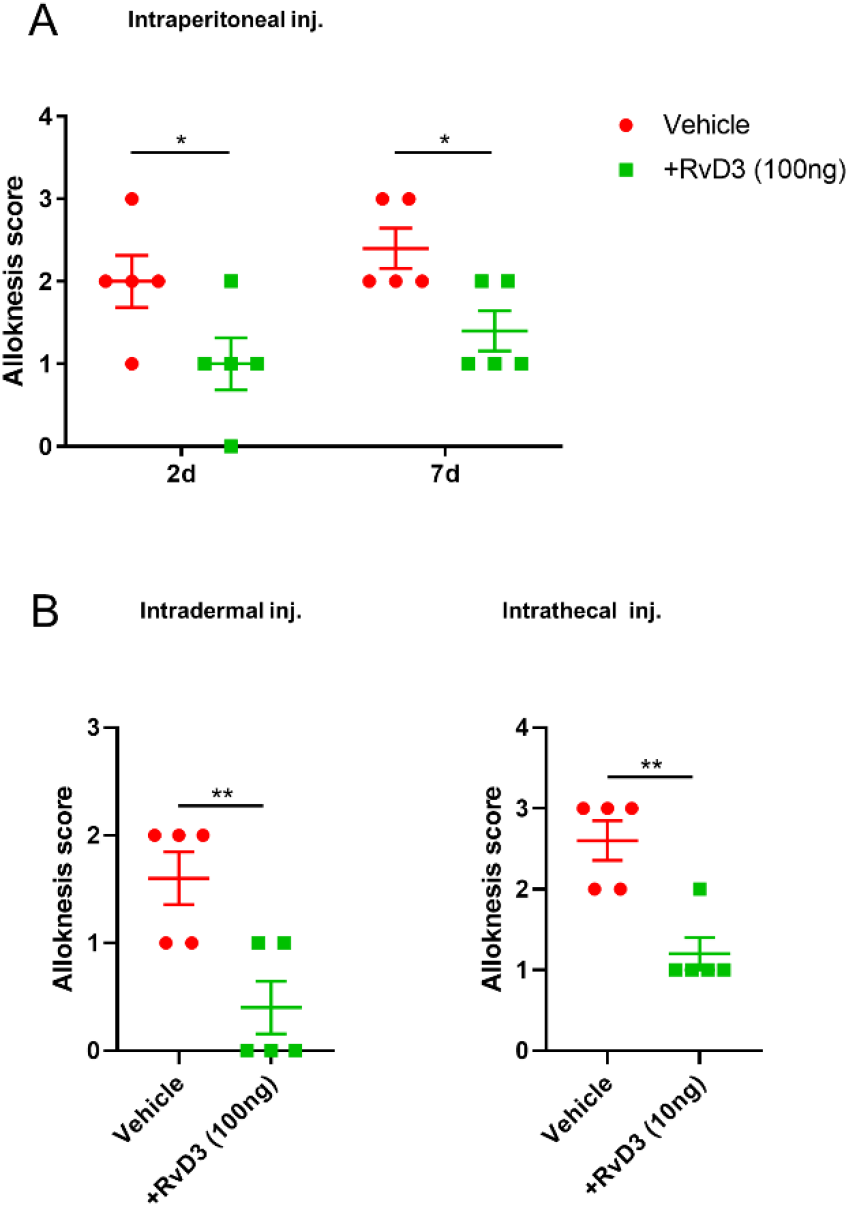
Effects of different routes of a single injection of RvD3 on psoriasiform alloknesis. **A**, Intraperitoneal single injection of RvD3 attenuated alloknesis on day 2 and 7 (n = 5 mice/group). **B**, Similar to intraperitoneal single injection, Intradermal and intrathecal single injection of RvD3 attenuated alloknesis on day 7 (n = 5 mice/group). Statistical analysis: (A) Two-way ANOVA, Bonferroni post-test. (B) Student’s t-test. Error bars indicate mean ± s.e.m. and *p < 0.05, **p < 0.01.

**Fig S2:**
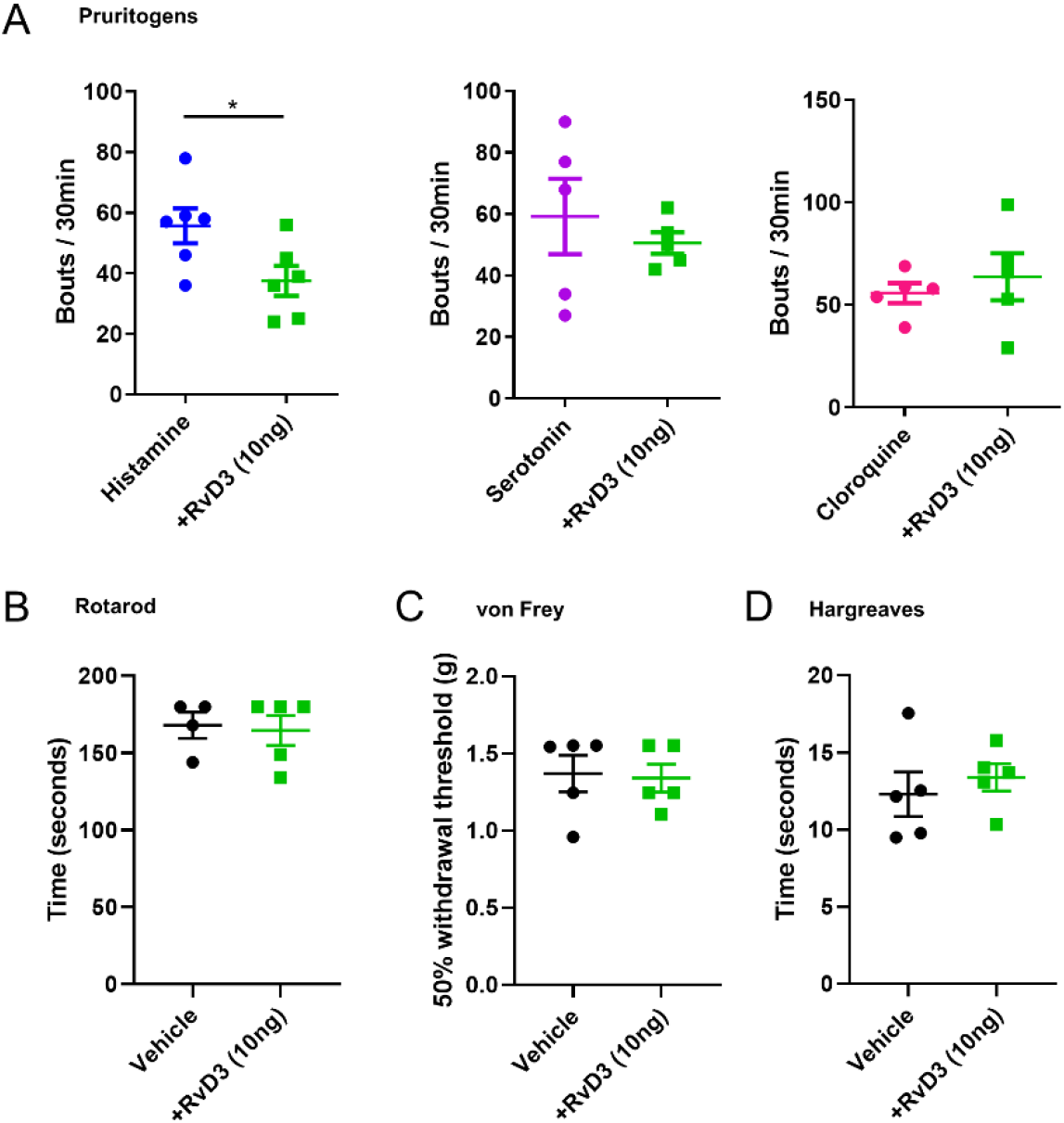
Effects of RvD3 on acute pain and itch. **A**, intradermal injection of RvD3 into the cheek inhibited histamine-provoked scratching, but not serotonin and chloroquine-provoked scratching (n = 5-6 mice/group). **B**, **C**, and **D**, Rotarod, von frey, and Hargreaves test are indistinguishable in vehicle- and RvD3-treated mice with intradermal injection into paw (n = 4-5 mice/group) Statistical analysis: (A, B, C, and D) Student’s t-test. Error bars indicate mean ± s.e.m. and *p < 0.05.

**Fig S3:**
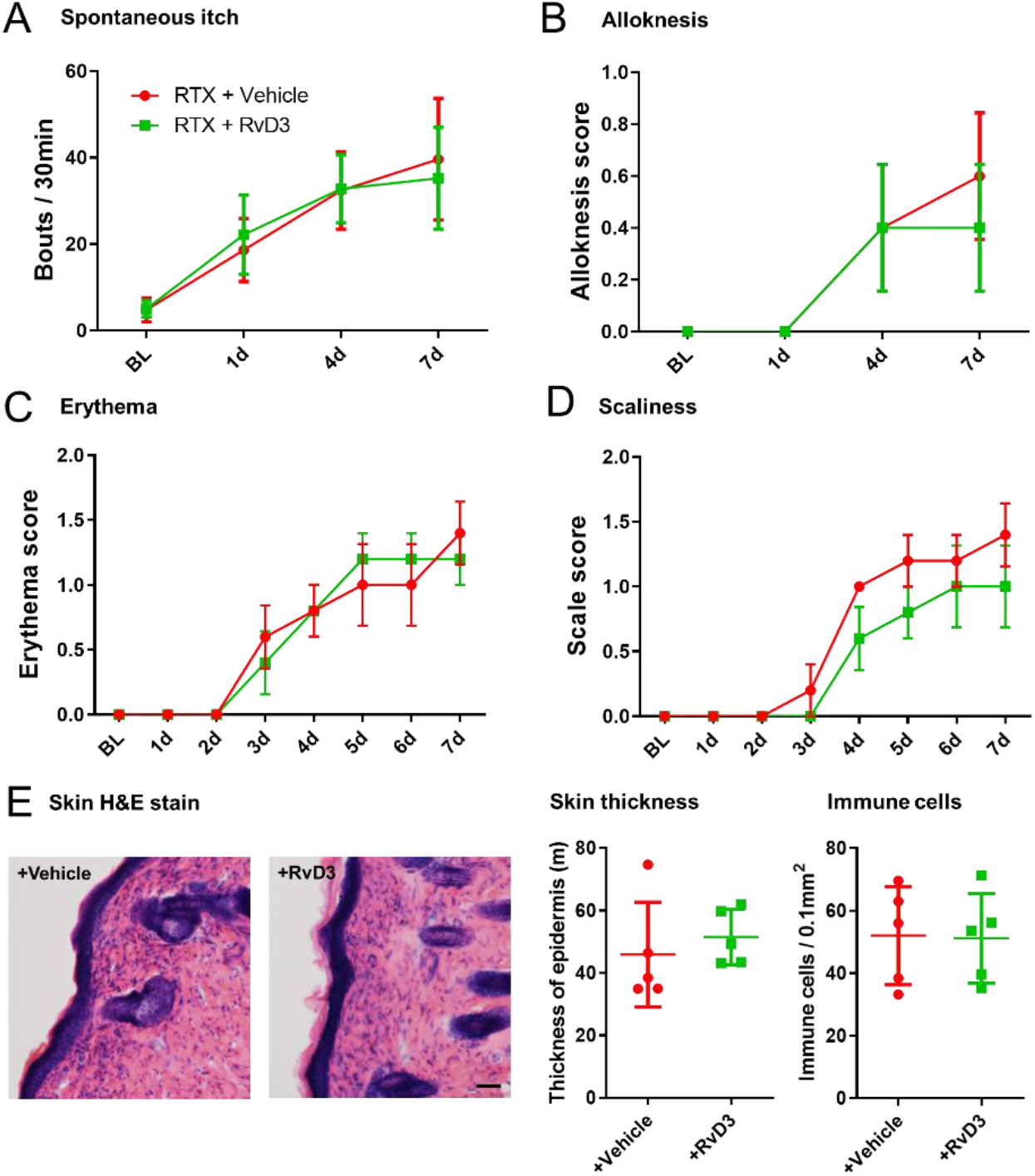
Effects of RTX and RvD3 treatments on psoriasis. **A** and **B**, Spontaneous itch and alloknesis of RTX-treated psoriasis model were not inhibited by repeated intraperitoneal injection of RvD3 (n = 5 mice/group). **C** and **D**, Erythema and scale score of skin lesion in RTX-treated psoriasis model were not regulated by RvD3 (n = 5 mice/group). **E**, Histopathology of skin lesion of RTX-treated psoriasis model with vehicle or RvD3 treatment. Scale bar, 50mm. Thickness of epidermis and number of immune cells were not altered by RvD3 treatment (n = 5 mice/group). Statistical analysis: (A, B, C, and D) Two-way ANOVA, Bonferroni post-test. (E) Student’s t-test. Error bars indicate mean ± s.e.m.

**Fig S4:**
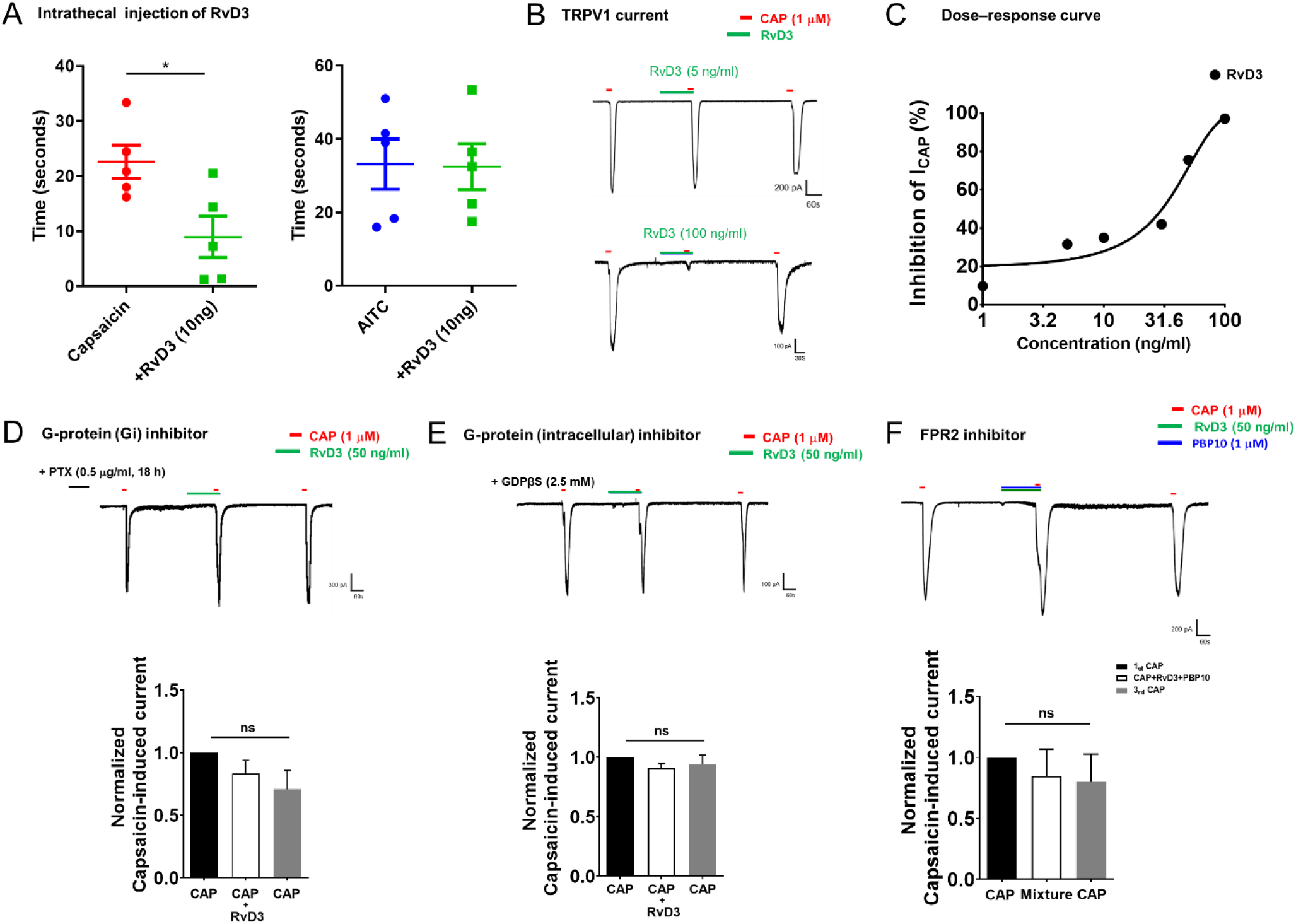
RvD3 inhibit TRPV1 activation via FPR2. **A**, Intrathecal injection of RvD3 inhibited capsaicin-induced acute pain, but not AITC-induced acute pain (n = 5 mice/group). **B**, Capsaicin-induced inward currents. Note a dose-dependent inhibition of TRPC1 currents by RvD3. **C**, Dose-response curves showing inhibition of TRPV1 currents by RvD3 (n = 4-6 neurons over 2mice). **D**, Pretreatment of DRG cultures with PTX blocks the inhibitory effects of RvD3 on TRPV1 current (n = 5 neurons over 2 mice). **E**, Intracellular perfusion of GDPbs (8 min) blocks the inhibitory effects of RvD3 on TRPV1 current (n = 5 neurons over 2 mice). **F**, FPR2 inhibitor (PBP10) blocks inhibitory effects of RvD3 on TRPV1 current (n = 5 neurons over 2 mice). Statistical analysis: (A) Student’s t-test. (D, E, and F) one-way ANOVA, Tukey post-tests. Error bars indicate mean ± s.e.m. and ns, not significant. *p < 0.05.

**Fig S5:**
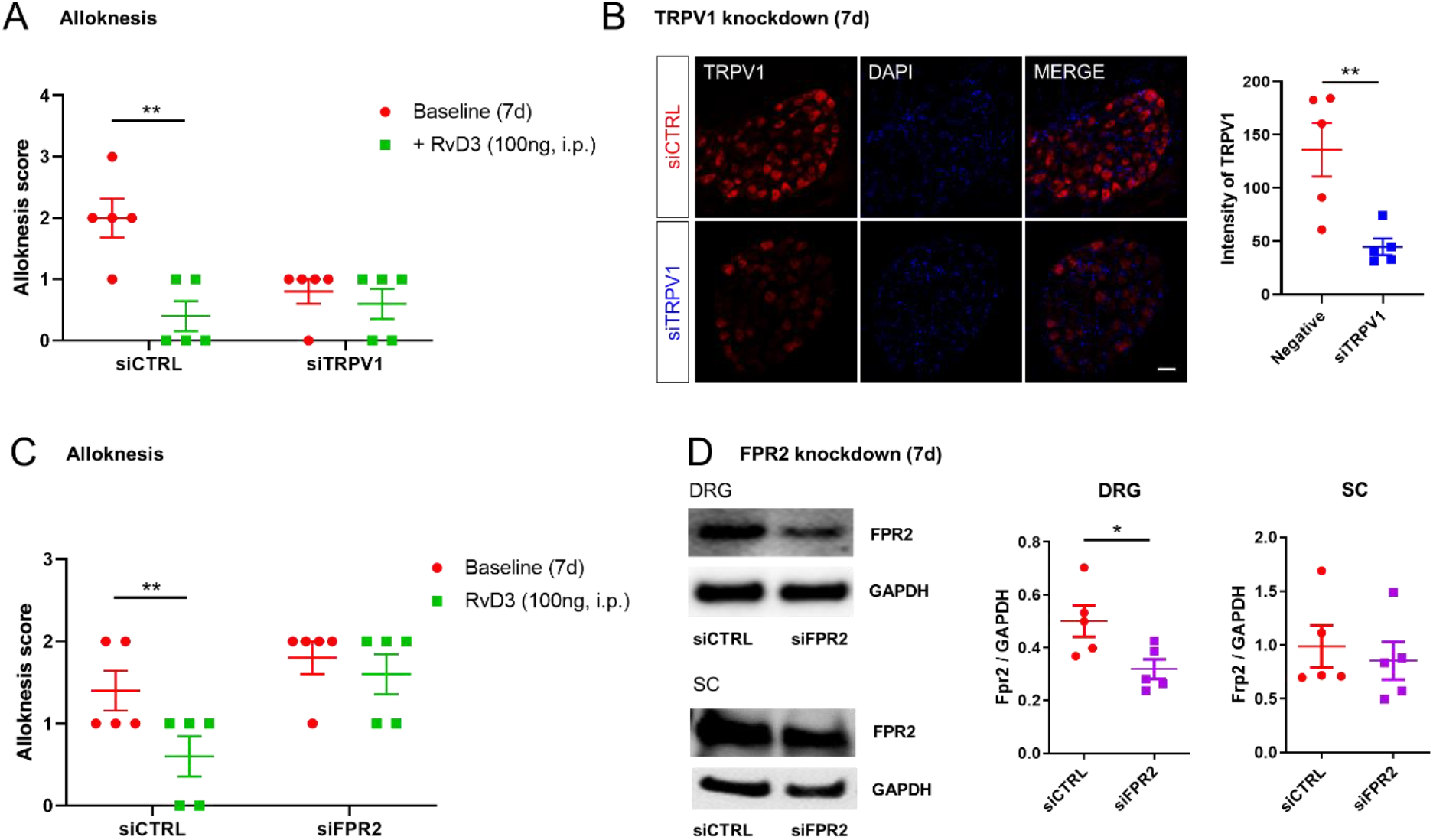
Effects of siTRPV1 and siFPR2 treatments on psoriasiform alloknesis. **A**, Intraperitoneal single injection of RvD3 didn’t inhibit alloknesis of siTRPV1-injected psoriasis model (n = 5 mice/group). **B**, Immunohistochemistry showing that siTRPV1 significantly reduced expression of TRPV1 in DRG (Scale bar, 50mm, n = 5 mice/group). **C**, Intraperitoneal single injection of RvD3 didn’t inhibit alloknesis of siFPR2-injected psoriasis model (n = 5 mice/group). **D**, Western blotting showing knockdown of FPR2 expression of approximately 40% to 50% in DRGs by FPR2 siRNA treatment, but not in spinal cord (n = 5 mice/group). Statistical analysis: (A and C) Two-way ANOVA, Bonferroni post-test. (B abd D) Student’s t-test. Error bars indicate mean ± s.e.m. and *p < 0.05, **p < 0.01.

**Fig S6:**
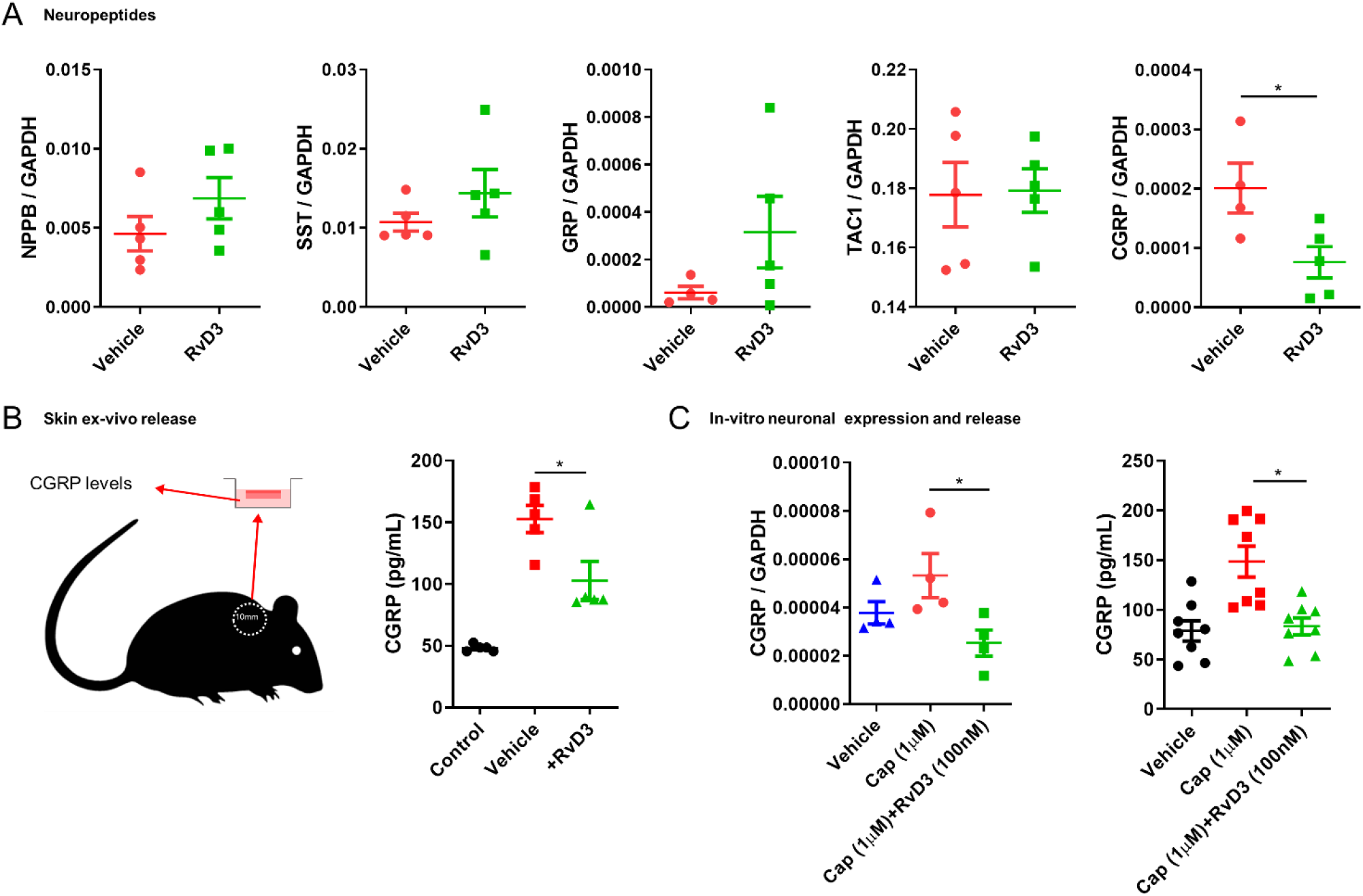
RvD3 regulated CGRP release in DRG neurons and psoriasiform skin. **A**, Transcriptional expression levels assessed by qPCR of neuropeptides (NPPB; natriuretic peptide type **B**, SST; somatostatin, GRP; gastrin releasing peptide, TAC1; tachykinin 1, CGRP; calcitonin gene-related peptide). Only CGRP in DRG of psoriasis model was downregulated by repeated intraperitoneal injection of RvD3 (n = 5 mice/group). B, Experimental design (skin explant): measurement of CGRP release ex vivo from nape skin lesion punch biopsies. Repeated intraperitoneal injection of RvD3 significantly reduced CGRP release form nape skin of psoriasis model. Samples of A and B were collected on day 7 of psoriasis model (n = 5 mice/group). **C**, Capsaicin-induced CGRP release from cultured DRG neurons was significantly inhibited by RvD3 (left; qPCR, n = 4, right; ELISA, n = 8). Statistical analysis: (A) Student’s t-test. (B and C) one-way ANOVA, Tukey post-tests. Error bars indicate mean ± s.e.m. and *p < 0.05.

**Fig S7:**
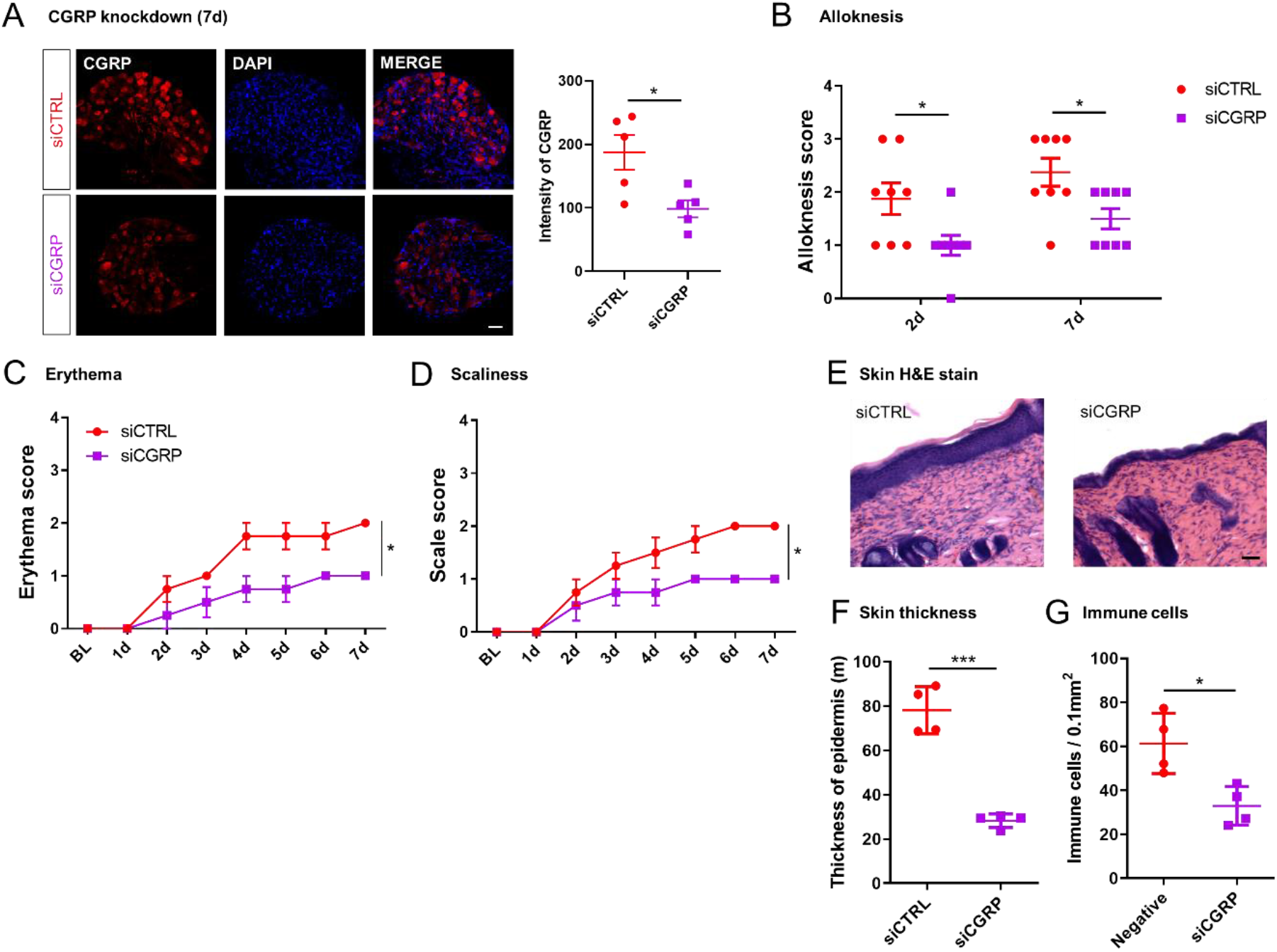
siCGRP attenuates psoriasiform itch and skin inflammation. **A**, Immunohistochemistry showing that siCGRP significantly reduced expression of TRPV1 in DRG (Scale bar, 50mm, n = 5 mice/group) **B**, Alloknesis of psoriasis model was significantly decreased in mice injected with CGRP siRNA (n = 8 mice/group). **C** and **D**, siCGRP significantly inhibited erythema and scaliness in psoriasis model (n = 4 mice/group). **E**, Histopathology of skin lesion from siCTRL and siCGRP-treated mice on day 7 of psoriasis model (Scale bar, 50mm). **F** and **G**, Thickness of epidermis and number of immune cells are significantly reduced in skin of siCGRP-treated psoriasis model (n = 4 mice/group). Statistical analysis: (A, F, and G) Student’s t-test. (B, C, and D) Two-way ANOVA, Bonferroni post-test. Error bars indicate mean ± s.e.m. and *p < 0.05, ***p < 0.01.

**Fig S8:**
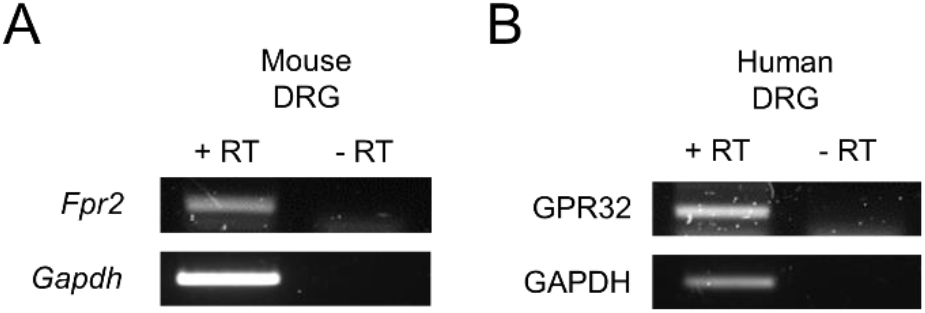
Expression of RvD3 receptors in mouse and human DRG tissue. **A**, PCR amplification of Fpr2 transcripts showed expression in mouse DRGs. **B**, PCR amplification of GPR32 transcripts showed expression in human DRGs. Experimental details and primer sequences are described in the Methods section.

